# Co-expression of IBDV VP2 and H9N2 HA by recombinant HVT induces high protection against both pathogens in chickens

**DOI:** 10.64898/2026.05.12.724538

**Authors:** Yuanyuan Zhang, Xiuwen Yang, Yunzhe Kang, Wenhui Zhu, Yuanyuan Sun, Shaoyan Qi, Yuxin Chen, Guoqing Zhuang, Aijun Sun

**Author notes:** Address correspondence to Guoqing Zhuang, Address correspondence to Aijun Sun. Yuanyuan Zhang and Xiuwen Yang contributed equally to this article. Author order was determined based on contribution to the article. Contributed equally.

## Abstract

Infectious bursal disease virus (IBDV) and H9N2 avian influenza virus (AIV) are significant global threats to poultry health and production. While IBDV induces severe immunosuppression, undermining host defense and vaccine efficacy, H9N2 AIV is characterized by widespread prevalence, persistent shedding, and substantial economic losses. Conventional inactivated vaccines often fail to elicit robust cellular immunity and necessitate multiple booster doses, underscoring the urgent requirement for advanced multivalent vaccination platforms. To address this, we developed a recombinant herpesvirus of turkey (rHVT BAC-VP2-HA) using a bacterial artificial chromosome (BAC) vector system, engineered to co-express the major protective antigen *VP2* of IBDV and the hemagglutinin (*HA*) of H9N2 AIV. Genetic stability and *in vitro* characterization confirmed that the recombinant exhibited replication kinetics and plaque morphology comparable to parental HVT, with stable antigen expression. In SPF chickens, rHVT BAC-VP2-HA induced strong humoral immune responses against both target antigens, comparable to those elicited by a commercial inactivated vaccine. Crucially, the recombinant virus significantly enhanced cellular immunity, evidenced by markedly elevated CD3^+^CD8^+^ T cell responses. Upon challenge, the recombinant conferred high clinical protection (86%) against virulent IBDV, significantly ameliorating bursal pathology and reducing viral loads. Notably, it provided complete (100%) protection against H9N2 AIV, effectively abolishing viral shedding and suppressing viral replication in respiratory tissues. These results demonstrate that rHVT BAC-VP2-HA is a safe and efficacious candidate capable of eliciting humoral and cellular immune responses, offering a promising strategy for the integrated control of major poultry diseases.

**Importance:** Infectious bursal disease virus (IBDV) and H9N2 avian influenza virus (AIV) are major pathogens that frequently co-circulate in poultry, where IBDV-induced immunosuppression compromises the efficacy of vaccination against other infectious diseases. Conventional inactivated vaccines primarily induce humoral immunity and are often insufficient to prevent viral shedding or provide broad protection against multiple pathogens. In this study, we developed a recombinant herpesvirus of turkeys (HVT) vaccine co-expressing the IBDV VP2 and H9N2 HA antigens and demonstrated that it induces both robust antibody responses and enhanced CD8^+^ T cell immunity. Notably, this vaccine not only provided effective protection against IBDV but also completely prevented viral shedding following H9N2 challenge. These findings highlight the advantage of HVT-vectored multivalent vaccines in eliciting balanced immune responses and controlling virus transmission, providing important insights for the development of next-generation vaccines against immunosuppressive and respiratory viral co-infections in poultry.

## 1. Introduction

Infectious bursal disease virus (IBDV) represents a major threat to the global poultry industry (Wang et al., 2021; Zhang et al., 2022). This double-stranded RNA virus, characterized by a bisegmented genome (segments A and B), facilitates genetic reassortment, driving the continuous emergence of variant strains with altered antigenicity (Fan et al., 2025; Zhang et al., 2026b). This rapid evolution promotes immune escape and compromises the efficacy of existing vaccines (Jing et al., 2026; Yu et al., 2025; Zhang et al., 2022). Infection results in infectious bursal disease (IBD), which is characterized by bursal atrophy, splenomegaly, and severe immunosuppression. Central to this pathology is VP2, the major capsid protein containing multiple conformational neutralizing epitopes (Guo et al., 2021; Zhang et al., 2025). Because genetic variation in VP2 is closely associated with antigenic shifts and immune escape, it is widely recognized as the primary protective antigen and an ideal target for vaccine development (Xiong et al., 2025).

Compounding these challenges, the avian influenza virus (AIV) poses a concurrent and significant threat to the poultry industry (Sun et al., 2023; Yang et al., 2023). AIV causes avian influenza (AI), an acute, highly contagious disease responsible for respiratory distress, decreased egg production, and substantial economic losses (Bruno et al., 2026; Kim et al., 2026). While AIVs are classified into subtypes based on hemagglutinin (HA) and neuraminidase (NA) surface glycoproteins, the H9N2 subtype has become particularly prevalent (Li et al., 2026; Liang et al., 2024; Luczo and Spackman, 2024). HA serves as the primary antigen for inducing neutralizing antibody-mediated protection. Crucially, the immunosuppression induced by IBDV compromises the host’s immune response, thereby reducing the protective efficacy of AIV vaccines. Consequently, the co-infection of IBDV and AIV represents a complex challenge for effective poultry health management (Spackman et al., 2018).

Vaccination remains one of the most economical and effective strategies for controlling IBD and AI. Currently, prevention mainly relies on conventional vaccines, including inactivated and live attenuated vaccines (Alqazlan et al., 2022; Wang et al., 2025; Xie et al., 2025). However, these vaccines have several limitations, including the risk of reversion to virulence, poor antigenic matching with circulating strains, and reduced efficacy against rapidly evolving viruses. Consequently, there is an urgent imperative to develop innovative vaccination strategies. In this context, viral vector-based platforms have emerged as a highly promising alternative (Hein et al., 2021).

Among the various viral vectors explored, Marek’s disease virus (MDV) has garnered significant attention due to its favorable biological properties. Specifically, herpesvirus of turkeys (HVT), a non-oncogenic serotype of MDV, has been established as a safe and efficacious vaccine vector (Ingrao et al., 2017; Wu et al., 2026). Recombinant HVT expressing heterologous antigens has demonstrated success in conferring protection against specific pathogens. For example, the insertion of the IBDV *VP2* gene into the intergenic region between *UL55* and *UL56* has yielded recombinant HVT capable of providing robust protection against virulent IBDV (vIBDV) strains (Shah et al., 2022). Similarly, HVT-vectored vaccines expressing H9N2 HA have proven effective in reducing viral shedding following H9N2 AIV challenge, primarily through the induction of potent mucosal immunity (Liu et al., 2019; Yang et al., 2025; Zai et al., 2022). These findings collectively underscore the potential of HVT-based vaccines to elicit protective immunity against both IBDV and H9N2 AIV.

However, a critical knowledge gap remains regarding the efficacy of a single HVT vector co-expressing both VP2 and HA. Specifically, it is unclear whether such a construct can simultaneously confer effective protection against both pathogens, induce balanced humoral and cellular immune responses, and effectively mitigate viral shedding.

To address this gap, an HVT (rHVT BAC-VP2-HA) co-expressing the IBDV VP2 and H9N2 AIV HA proteins was constructed. The *VP2* gene was inserted into the *UL55* locus, while the *HA* gene was inserted into the *RR* locus. The recombinant virus was characterized *in vitro* for growth kinetics and genetic stability, and subsequently evaluated *in vivo* for immunogenicity and protective efficacy in SPF chickens. Notably, rHVT BAC-VP2-HA elicited robust humoral immune responses comparable to those induced by a commercial inactivated vaccine, while simultaneously stimulating enhanced cellular immunity, particularly the activation of CD8^+^ T cells. Upon challenge, the recombinant vaccine provided effective protection against vIBDV. Importantly, it completely prevented viral shedding following H9N2 AIV infection. These results indicate that rHVT BAC-VP2-HA is a highly promising multivalent vaccine candidate, offering the dual advantage of conferring comprehensive protection and limiting virus transmission.

## 2. Materials and methods

### 2.1 Viruses, cells, and plasmids

H9N2 virus strain A/chicken/Henan/Zk2/2012 was propagated in 9-day-old specific pathogen-free (SPF) embryonated chicken eggs, while virulent IBDV was propagated in the DF-1 chicken embryonic fibroblast cell line. The recombinant virus (rHVT BAC-VP2-HA) was propagated in primary chicken embryonic fibroblasts (CEFs). Virus titers were determined as 50% egg infectious doses (EID_50_) for H9N2 or plaque-forming units (PFU) for cell culture-propagated viruses. CEFs were prepared from 9-day-old SPF chicken embryos (Boehringer Ingelheim, Beijing, China). CEFs and DF-1 cells were maintained in Dulbecco’s Modified Eagle’s Medium (DMEM; Sigma-Aldrich, USA) supplemented with 5% or 10% fetal bovine serum (FBS; Gibco, USA), respectively, and incubated at 37°C with 5% CO_2_. The HVT BAC infectious clone and shuttle plasmids pcDNA3.1-*VP2*-*kan* and pCAGGS-*HA*-*kan* were maintained in our laboratory.

### 2.2 Construction of recombinant rHVT BAC-VP2-HA infectious clone

To generate the recombinant rHVT BAC-VP2-HA infectious clone, a two-step Red-mediated recombination strategy was employed, as previously described (Tischer and Kaufer, 2012; Tischer et al., 2006). The HVT BAC backbone was maintained in recombination-competent *Escherichia coli (E. coli)*. In the first recombination step, the CMV-VP2-kan^R^-polyA expression cassette was amplified from the shuttle plasmid pcDNA3.1-VP2-kan^R^ using primers (VP2-kan^R^-F and VP2-kan_R_-R; Table 1) that contained homologous arms targeting the *UL55* locus. The purified PCR product was electroporated into *E. coli* harboring the HVT BAC, and recombinant clones were selected on kanamycin-containing medium. Correct insertion at the target locus was confirmed by PCR. In the second step, expression of the I-SceI endonuclease was induced by L-arabinose, leading to cleavage within the inserted cassette and promoting a second homologous recombination event that excised the kan_R_ selection marker. Successful deletion of the *kan* gene was confirmed by PCR, and the resulting rHVT BAC-VP2 clone was further validated by DNA sequencing to exclude unintended mutations. Subsequently, the rHVT BAC-VP2-HA clone was generated using the same strategy. The *HA-kan* cassette was inserted into the *RR* locus of rHVT BAC-VP2, followed by I-SceI-mediated excision of the kan^R^ selection marker.

**Table 1.**
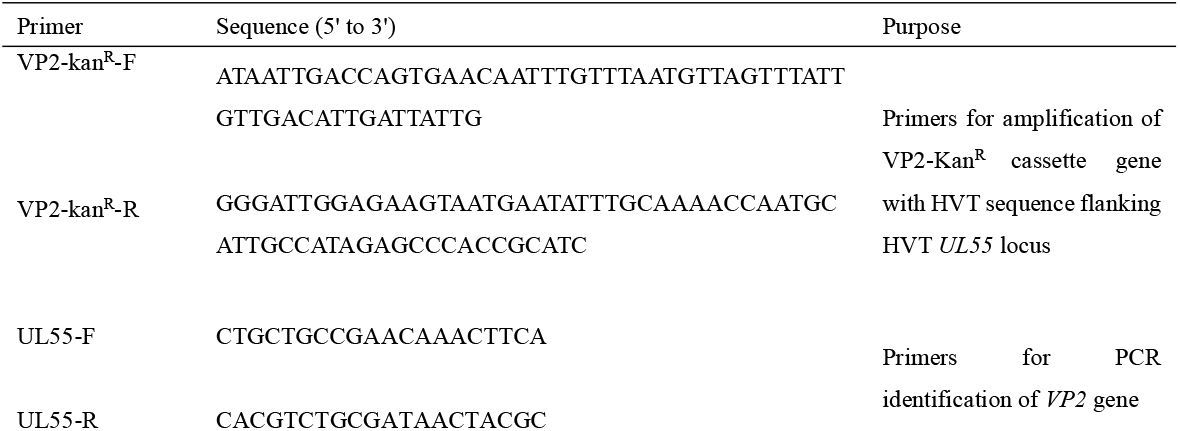
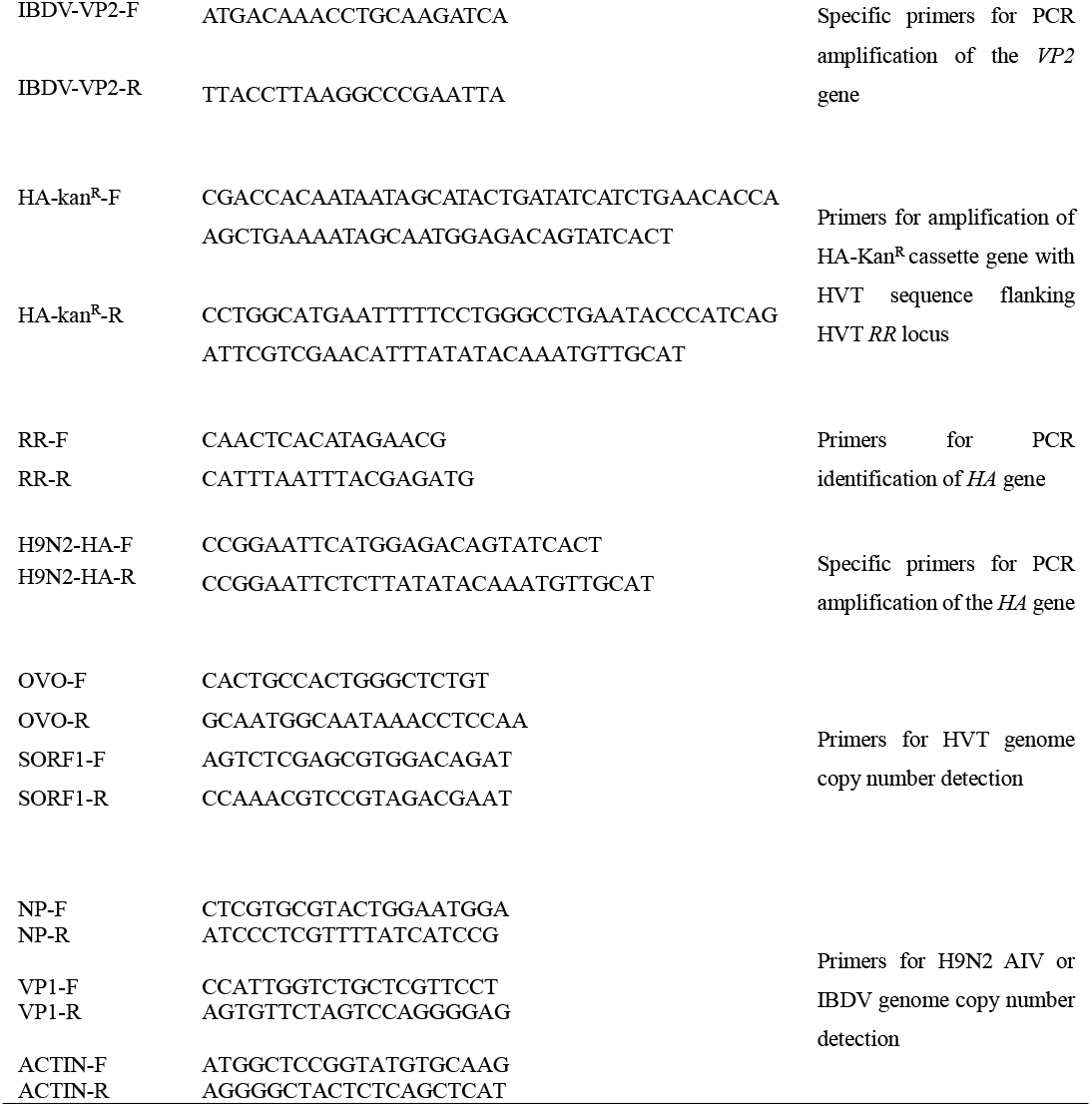
List of primers used in the study.

### 2.3 Rescue and identification of rHVT BAC-VP2-HA

The recombinant plasmid rHVT BAC-VP2-HA (500 ng per 60-mm dish) was transfected into CEFs using Lipofectamine™ 2000 (Invitrogen, USA) following the manufacturer’s protocols. Briefly, CEFs at 70-80% confluence were incubated with DNA-lipid complexes prepared in Opti-MEM. After 6 h, the transfection medium was replaced with maintenance medium containing 1% FBS and penicillin-streptomycin. Cells were cultured until cytopathic effects (CPE) appeared. Once CPE reached approximately 60%, infected cells were harvested and serially passaged onto fresh CEF monolayers in 35-mm dishes for 3-5 days to amplify the recombinant virus. Cells exhibiting approximately 60% CPE were subsequently fixed with 4% paraformaldehyde for further analyses.

Expression of HVT, IBDV VP2, and H9N2 AIV HA antigens was confirmed by indirect immunofluorescence assay (IFA). For detection, HVT polyclonal antisera (1:50) and H9N2 polyclonal antisera (1:50) were used as primary antibody, followed by FITC-conjugated goat anti-chicken IgG as the secondary antibody. Monoclonal antibody against IBDV VP2 protein (1:2000) was used as primary antibody, followed by FITC-conjugated goat anti-mouse IgG. In parallel, viral genomic DNA was extracted from infected cells and subjected to PCR and sequencing to verify the correct insertion and integrity of the foreign gene cassette.

### 2.4 Biological characterization of rHVT BAC-VP2-HA *in vitro*

#### 2.4.1 Growth kinetics assay

To evaluate the replication capacity of the recombinant virus rHVT BAC-VP2-HA, confluent monolayers of CEFs in six-well plates were separately infected with 100 PFU of either rHVT BAC-VP2-HA or the parental HVT BAC. Infected cells were harvested at 24-hour intervals from 24 to 120 hours post-infection (hpi). Viral genomic DNA was extracted using the phenol-chloroform method, and DNA concentrations were measured using a NanoDrop ND-1000 spectrophotometer (Thermo Fisher Scientific; MA, USA). All DNA samples were subsequently normalized to 10 ng/μL prior to downstream analysis.

Quantitative real-time PCR (qPCR) was performed using SYBR Green Master Mix (Thermo Fisher Scientific, USA) on a QuantStudio 5 system (Applied Biosystems, USA). Each 20 μL reaction mixture contained 10 μL of SYBR Green Master Mix, 0.5 μM of each primer, and approximately 50 ng of template DNA. The thermal cycling protocol consisted of an initial denaturation step at 95°C for 10 min, followed by 40 cycles of 95°C for 15 s and 60°C for 60 s. Melting curve analysis was subsequently conducted to confirm amplification specificity.

Standard curves were generated using serial dilutions of the recombinant HVT BAC plasmid and the pMD18-OVO plasmid, yielding the following regression equations: for *SORF1*, y = –3.5147x + 37.778 (R^2^ = 0.9989); and for *OVO*, y = –3.2612x + 24.091 (R^2^ = 0.9983). Viral genome copy numbers were calculated based on standard curves using primers specific for *SORF1* (targeting the HVT genome) and *OVO* (serving as an endogenous reference gene for CEFs). Viral loads were normalized to the *OVO* gene and expressed as the *SORF1*/*OVO* copy number ratio, representing the number of viral genome copies per cell equivalent. Growth curves and statistical analyses were performed using GraphPad Prism 8.

#### 2.4.2 Plaque size determination

To compare plaque morphology, confluent monolayers of CEFs in 35-mm dishes were inoculated with 100 PFU of either the recombinant rHVT BAC-VP2-HA or the parental HVT BAC. At 5 days post-infection (dpi), when distinct plaques became visible, the cells were washed with PBS and fixed with ice-cold acetone/methanol (3:2, v/v) for 5 min. After air-drying, the cells were blocked with 5% (w/v) skim milk in PBS at 37°C for 1 h, followed by three washes with PBS (5 min each) on a shaker.

For immunostaining, the fixed cells were incubated with HVT polyclonal antiserum at 37°C for 1 h. After three additional PBS washes, a goat anti-chicken FITC-conjugated secondary antibody was applied and incubated at 37°C for 1 h in the dark. Following a final PBS wash, plaques were visualized using an inverted fluorescence microscope (Olympus, Japan). For each virus, the areas of 50 randomly selected plaques were measured using ImageJ. Relative plaque sizes were calculated by normalizing to the mean plaque area of the parental HVT BAC, which was set as 100%. All experiments were performed in triplicate as three independent biological replicates.

#### 2.4.3 Recombinant virus genetic stability assay

To assess the genetic stability of the recombinant virus rHVT BAC-VP2-HA during serial propagation, confluent monolayers of CEFs in 60-mm dishes were inoculated with 100 PFU of the virus. Infected cells were harvested every 3 days and subsequently passaged onto fresh CEF monolayers for continuous propagation which was maintained for up to 20 serial passages.

At every fifth passage, the expression of VP2 and HA proteins was verified by IFA to confirm the stable maintenance of the inserted genes.

### 2.5 Evaluation of the safety and immunogenicity of rHVT BAC-VP2-HA in SPF chickens

#### 2.5.1 Vaccination and experimental design

One-day-old SPF chickens were randomly assigned into three groups (n = 45 per group). The groups consisted of a mock group that received an equivalent volume of DMEM, a commercial inactivated vaccine group, and an rHVT BAC-VP2-HA group, in which each chick was subcutaneously inoculated with 2,000 PFU of the recombinant virus. All immunizations were administered via subcutaneous injection in the cervical region.

#### 2.5.2 Growth performance and clinical observation

To evaluate the impact of rHVT BAC-VP2-HA vaccination on growth performance, body weights were recorded weekly for five randomly selected chickens from each group over a 5-week period following vaccination. In parallel, all chickens were monitored daily for any clinical signs throughout the same period.

#### 2.5.3 Humoral immune responses

To evaluate humoral immune responses, blood samples were collected weekly from the wing veins of five randomly selected chickens per group over a 5-week period following vaccination. Serum was subsequently separated, and analyzed for HA- and VP2-specific antibody responses using a hemagglutination inhibition (HI) assay and an indirect enzyme-linked immunosorbent assay (ELISA), respectively.

#### 2.5.4 Cellular immune responses

To evaluate cellular immune responses, spleens were collected from five randomly selected chickens per group at 28 days post-vaccination. Single-cell suspensions were prepared using a commercial kit (Solarbio, China) following the manufacturer’s instructions, and the cells were subsequently resuspended in PBS at a concentration of 1.0 × 10^6^ cells/mL.

For surface staining, 1.0 × 10^6^ cells suspended in 1 mL PBS were incubated with a cocktail of fluorochrome-conjugated antibodies specific for chicken CD3 (AF647), CD4 (PB450), CD8a (FITC), and Bu-1 (PE) (Southern Biotech, AL, USA) for 30 min at 4°C in the dark. Cells were then washed three times with ice-cold PBS (300 × g, 5 min, 4°C) and resuspended in 500 μL PBS. Stained cells were then acquired on a flow cytometer, and the resulting data were analyzed to determine the frequencies of T and B lymphocyte subsets.

### 2.6 Evaluation of protective efficacy against IBDV and H9N2 AIV in SPF chickens

At 28 days post-vaccination, chickens from each group were randomly allocated into three subgroups of 15 chickens each (Table 2). Chickens in two of the subgroups were challenged via the oculonasal route with IBDV (10^4^ EID_50_/100 μL) or H9N2 AIV (10^4^ EID_50_/100 μL), while the remaining subgroup served as unchallenged controls.

**Table 2.** Experimental design of challenge study.

| Group | No. of chickens | Immunization (Day 1) | Challenge (Day 28) | No. of chickens |
| --- | --- | --- | --- | --- |
| Mock | 45 | DMEM | IBDV | 15 |
|  |  |  | H9N2 AIV | 15 |
|  |  |  | Unchallenged | 15 |
| Commercial<br>inactivated vaccine | 45 | Inactivated vaccine | IBDV | 15 |
|  |  |  | H9N2 AIV | 15 |
|  |  |  | Unchallenged | 15 |
| rHVT BAC-VP2-HA | 45 | rHVT BAC-VP2-HA | IBDV | 15 |
|  |  |  | H9N2 AIV | 15 |
|  |  |  | Unchallenged | 15 |

#### 2.6.1 Protective efficacy against IBDV challenge

Following challenge, clinical signs and mortality were monitored and recorded daily for 7 consecutive days. Chickens exhibiting severe clinical signs were humanely euthanized and all remaining birds were euthanized at 7 days post-challenge (dpc).

At 7 dpc, body weights were recorded prior to euthanasia. The bursae of Fabricius were collected and weighed. The bursa-to-body weight ratio (BBW) was calculated as (bursa weight/body weight) × 1000. The bursa body index (BBIX) was determined as the ratio of the BBW of the experimental group to that of the uninfected control group, with a BBIX value < 0.7 considered indicative of bursal atrophy.

For histopathological analysis, bursal tissues were fixed in 4% paraformaldehyde, embedded in paraffin, sectioned, and stained with hematoxylin and eosin (HE) for microscopic evaluation of pathological lesions.

For viral load determination, a portion of each bursa was homogenized, and total RNA was extracted using TRIzol reagent (Invitrogen, Carlsbad, CA, USA) according to the manufacturer’s instructions. Complementary DNA (cDNA) was synthesized using a High-Capacity cDNA Reverse Transcription Kit (Thermo Fisher Scientific, USA). qPCR was performed using VP1-specific primers (VP1-F/R), and viral genome copy numbers were determined based on standard curves.

#### 2.6.2 Protective efficacy against H9N2 challenge

Following challenge, clinical signs and mortality were monitored and recorded daily for 7 consecutive days. Chickens exhibiting severe clinical signs were humanely euthanized and all remaining birds were euthanized at 7 dpc.

To assess virus shedding, oropharyngeal and cloacal swabs were collected from each chicken at 3, 5, and 7 dpc. Swabs were eluted in antibiotic-supplemented PBS (penicillin 1,000 U/mL, streptomycin 1,000 μg/mL, and gentamicin 500 μg/mL), vortexed, and centrifuged at 2,000 rpm for 5 min at 4°C. Supernatants from oropharyngeal and cloacal swabs collected from the same chicken were pooled in equal volumes, filtered through a 0.22 μm membrane, and inoculated into the allantoic cavity of 9-day-old SPF embryonated chicken eggs. Embryos that died within 24 hours post-inoculation (hpi) were discarded. Allantoic fluids were harvested at 96 hpi and subjected to hemagglutination assay to detect infectious virus.

At 7 dpc, tracheal tissues were collected for histopathological and viral load analyses. For histopathological examination, tissues were fixed in 4% paraformaldehyde, embedded in paraffin, sectioned, and stained with HE for microscopic evaluation. For viral load determination, total RNA was extracted from tracheal tissues, reverse transcribed into cDNA, and subjected to qPCR using primers specific to H9N2 *NP* gene to quantify viral genome copy numbers.

### 2.7 Statistical analysis

All statistical analyses were performed using GraphPad Prism 8 (GraphPad Software, CA, USA). Data are presented as mean ± standard deviation (SD). Differences between two groups (including plaque size and viral growth kinetics) were analyzed using Student’s t-test. Comparisons among three or more groups were performed using one-way analysis of variance (ANOVA), followed by appropriate post hoc tests. A *P* < 0.05 was considered statistically significant.

## 3. Results

### 3.1 Successful construction and rescue of the rHVT expressing IBDV VP2 and H9N2 AIV HA proteins

Based on the infectious clone of HVT BAC, a recombinant HVT BAC expressing the IBDV *VP2* gene was first generated via a two-step Red-mediated homologous recombination strategy, with the *VP2* gene inserted into the *UL55* locus (Fig. 1A and B). Subsequently, the H9N2 AIV *HA* gene was inserted into the *RR* locus to construct rHVT BAC-VP2-HA (Fig. 1A and C).

**Figure 1.**
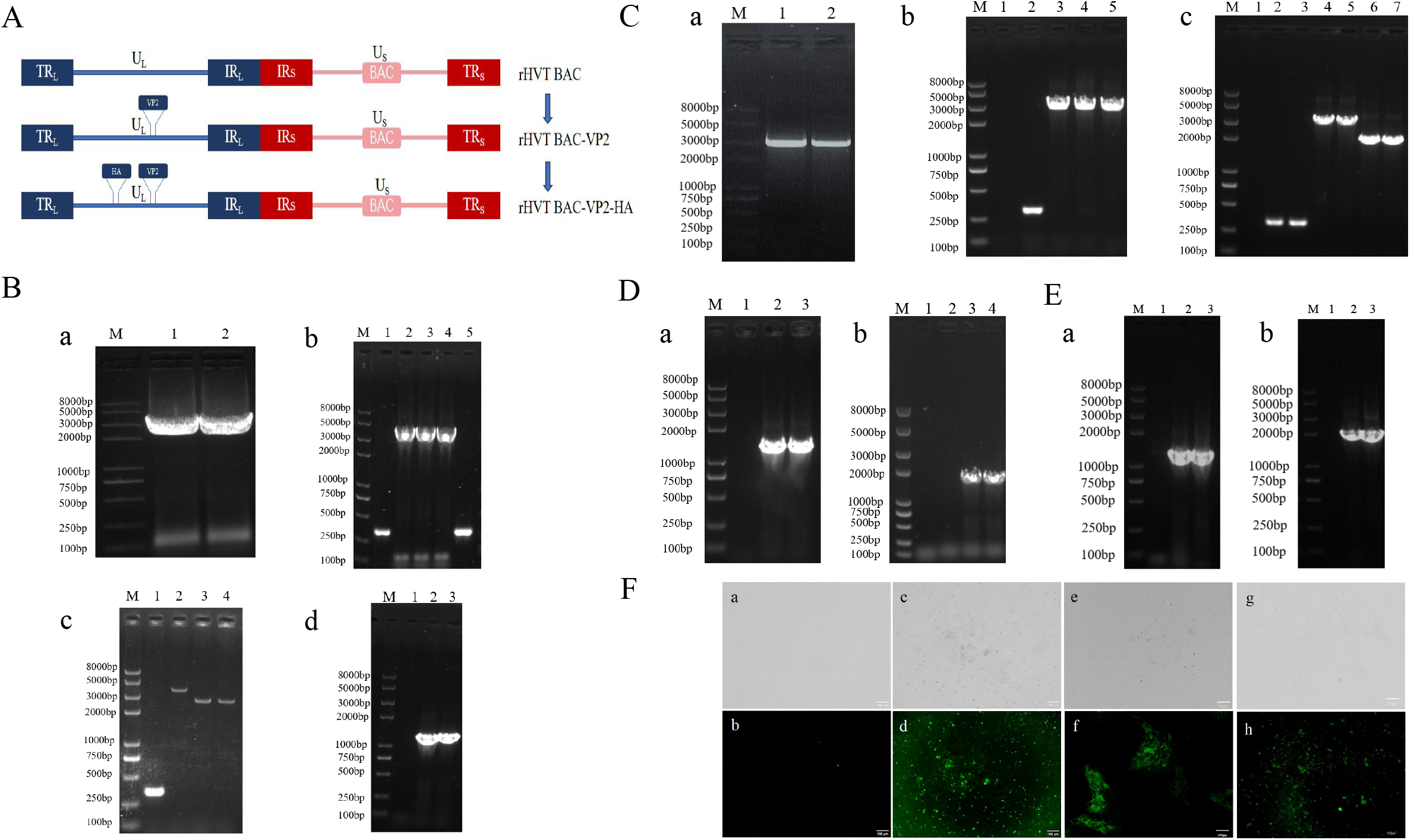
Construction and identification of the recombinant rHVT expressing VP2 and HA proteins. **(A)**. Schematic presentation of the construction of rHVT BAC-VP2-HA. HVT genome consists of a unique long (UL) region and a unique short (US) region, each flanked by inverted terminal and internal repeat long (TRL, IRL) and short (TRS, IRS) region. **(B)**. Construction of rHVT BAC-VP2 plasmid. (a). PCR amplification of the *VP2-kan* gene (expected band: 3430 bp). (b). PCR identification of the recombinant plasmid rHVT BAC-VP2-kan (expected band: 3739 bp). (c). PCR identification of the recombinant plasmid rHVT BAC-VP2 (expected band: 2689 bp). (d). PCR identification of the *VP2* gene in the recombinant plasmid rHVT BAC-VP2 (expected band: 1359 bp). DNA ladder sizes are indicated in bp. **(C)**. Construction of rHVT BAC-VP2-HA plasmid. (a). PCR amplification of the *HA-kan* gene (expected band: 3000 bp). (b). PCR identification of the recombinant plasmid rHVT BAC-VP2-HA-kan (expected band: 3100 bp). (c). PCR identification of the recombinant plasmid rHVT BAC-VP2-HA (expected band: 2000 bp). DNA ladder sizes are indicated in bp. **(D)**. PCR identification of foreign gene insertion in rHVT BAC-VP2-HA plasmid. (a). PCR identification of *HA* gene (expected band: 1683 bp). (b). PCR identification of *VP2* gene (expected band: 1359 bp). **(E)**. The presence of foreign genes in the recombinant viral DNA extracted from infected cells was verified by PCR. (a). PCR identification of *HA* gene (expected band: 1683 bp). (b). PCR identification of *VP2* gene (expected band: 1359 bp). **(F)**. Expression of HVT, IBDV VP2, and H9N2 AIV HA antigens was confirmed by indirect immunofluorescence assay (IFA). (a) and (b). CEF negative control. (c) and (d). Detection of HVT antigen using HVT polyclonal antiserum. (e) and (f). Detection of IBDV VP2 protein using a VP2-specific monoclonal antibody. (g) and (h). Detection of H9N2 AIV HA antigen using H9N2 polyclonal antiserum.

The successful construction of recombinant plasmids was confirmed by PCR analysis. Specific bands of the expected sizes were amplified using *VP2*- and *HA*-specific primers, respectively (Fig. 1D). Sequencing analysis further verified the correct insertion and sequence integrity of the foreign genes, demonstrating that *VP2* and *HA* were successfully integrated into the rHVT BAC genome. Collectively, these results confirm the successful construction of the recombinant plasmid rHVT BAC-VP2-HA.

To rescue the recombinant virus rHVT BAC-VP2-HA, CEF cells were transfected using the calcium phosphate precipitation method. Viral DNA extracted from infected cells was subjected to PCR analysis, which confirmed the presence of *VP2* and *HA* genes, indicating successful virus rescue (Fig. 1E). Expression of the foreign genes was further verified by IFA (Fig. 1F). Clear specific fluorescence signals were detected in infected CEF cells using HVT polyclonal antibody, IBDV VP2 monoclonal antibody, and H9N2 AIV polyclonal antibody, whereas no fluorescence signal was observed in the control group. Collectively, these results confirm the successful rescue of rHVT BAC-VP2-HA and the correct expression of both *VP2* and *HA* in infected cells.

### 3.2 The rHVT BAC-VP2-HA demonstrated excellent growth characteristics and genetic stability *in vitro*

To evaluate the effect of foreign gene insertion on viral replication, the plaque size and growth kinetics of rHVT BAC-VP2-HA were compared with those of the parental HVT BAC. Plaque assays revealed no significant difference in plaque area between rHVT BAC-VP2-HA and HVT BAC (*P* > 0.05; Fig. 2A). Consistently, growth curve analysis demonstrated comparable viral genome copy numbers between the two viruses at 1, 2, 3, 4, and 5 dpi (*P* > 0.05; Fig. 2B). These results indicate that the insertion of the VP2-HA cassette did not adversely affect viral replication *in vitro*. The genetic stability of rHVT BAC-VP2-HA was further assessed by serial passaging in CEFs for 20 passages.

**Figure 2.**
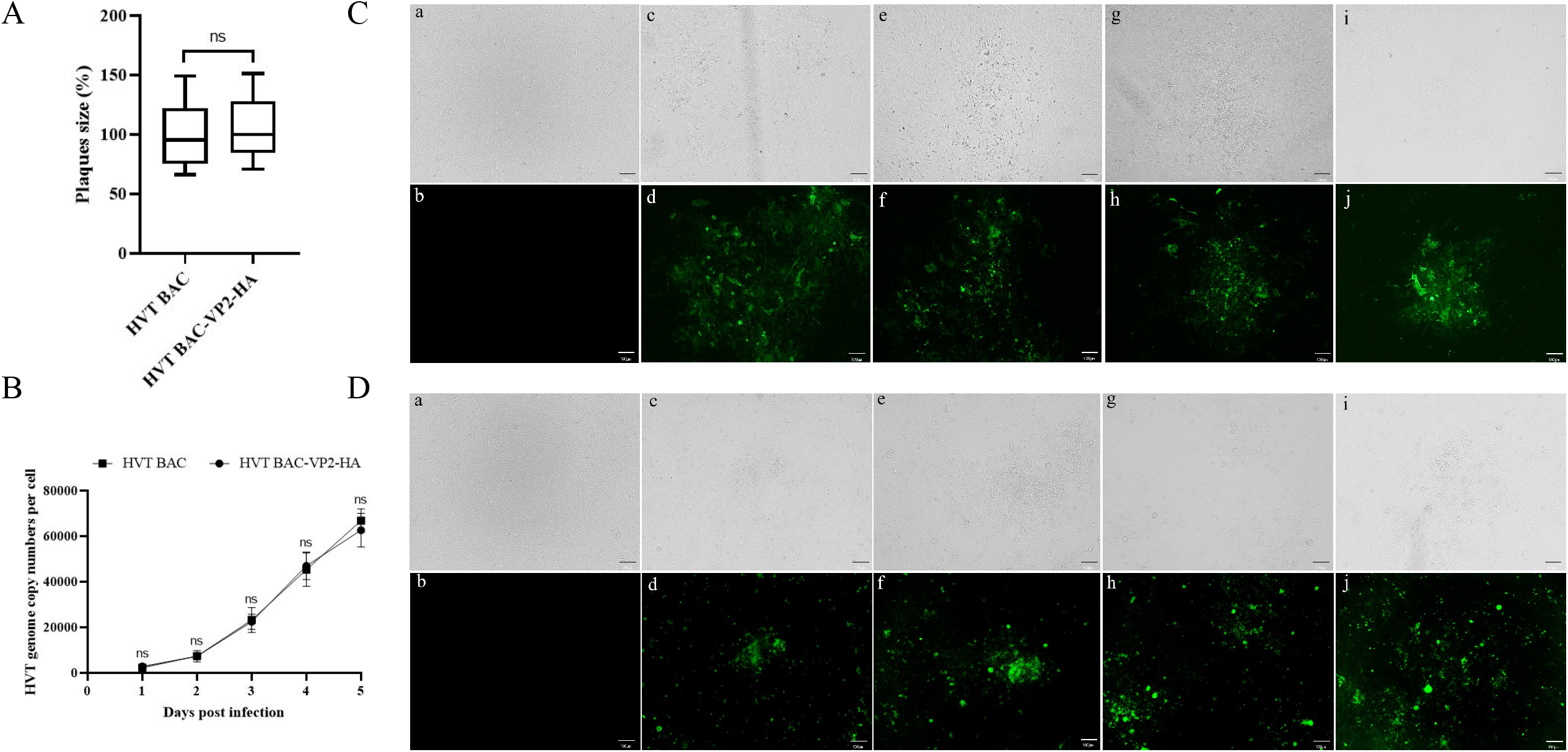
*In vitro* biological characterization of recombinant rHVT BAC-VP2-HA. **(A)**. The relative plaque size of the rHVT BAC-VP2-HA virus was determined at 7 dpi. The average plaque size of parental HVT BAC virus was set to 100%. **(B)**. *In vitro* growth kinetics. CEFs seeded on six-well plates were infected with 100 PFU of HVT BAC or rHVT BAC-VP2-HA viruses. At 1, 2, 3, 4, and 5 dpi, infected cells were harvested by trypsinization, and DNA was extracted for qPCR analysis to quantify HVT genome copy numbers. Each time point represents the mean value of triplicates in two independent experiments, with the error bar representing the standard error of the mean (SEM). **(C) and (D)**. Genetic stability of the VP2 and HA genes in the recombinant virus, detected using a monoclonal antibody against the IBDV VP2 protein and polyclonal antiserum against AIV H9N2, respectively. (a) and (b). CEF negative control. (c) and (d). CEFs infected with the 5th passage of the recombinant virus. (e) and (f). CEFs infected with the 10th passage of the recombinant virus. (g) and (h). CEFs infected with the 15th passage of the recombinant virus. (i) and (j). CEFs infected with the 20th passage of the recombinant virus.

The expression of VP2 and HA was examined every five passages by IFA. Specific fluorescence signals corresponding to VP2 and HA were consistently detected in infected cells across all passages, confirming stable retention and expression of the foreign genes throughout serial passaging.

### 3.3 rHVT BAC-VP2-HA induces robust humoral and cellular immune responses without affecting growth performance in SPF chickens

To evaluate the impact of vaccination on host growth, one-day-old SPF chickens were randomly divided into three groups and immunized with rHVT BAC-VP2-HA, a commercial inactivated vaccine, or DMEM as a mock control. Body weights were recorded at 7-day intervals post-immunization. All groups exhibited progressive weight gain over time, with no significant differences among the three groups throughout the experimental period (Fig. 3A), indicating that rHVT BAC-VP2-HA vaccination does not adversely affect growth performance.

**Figure 3.**
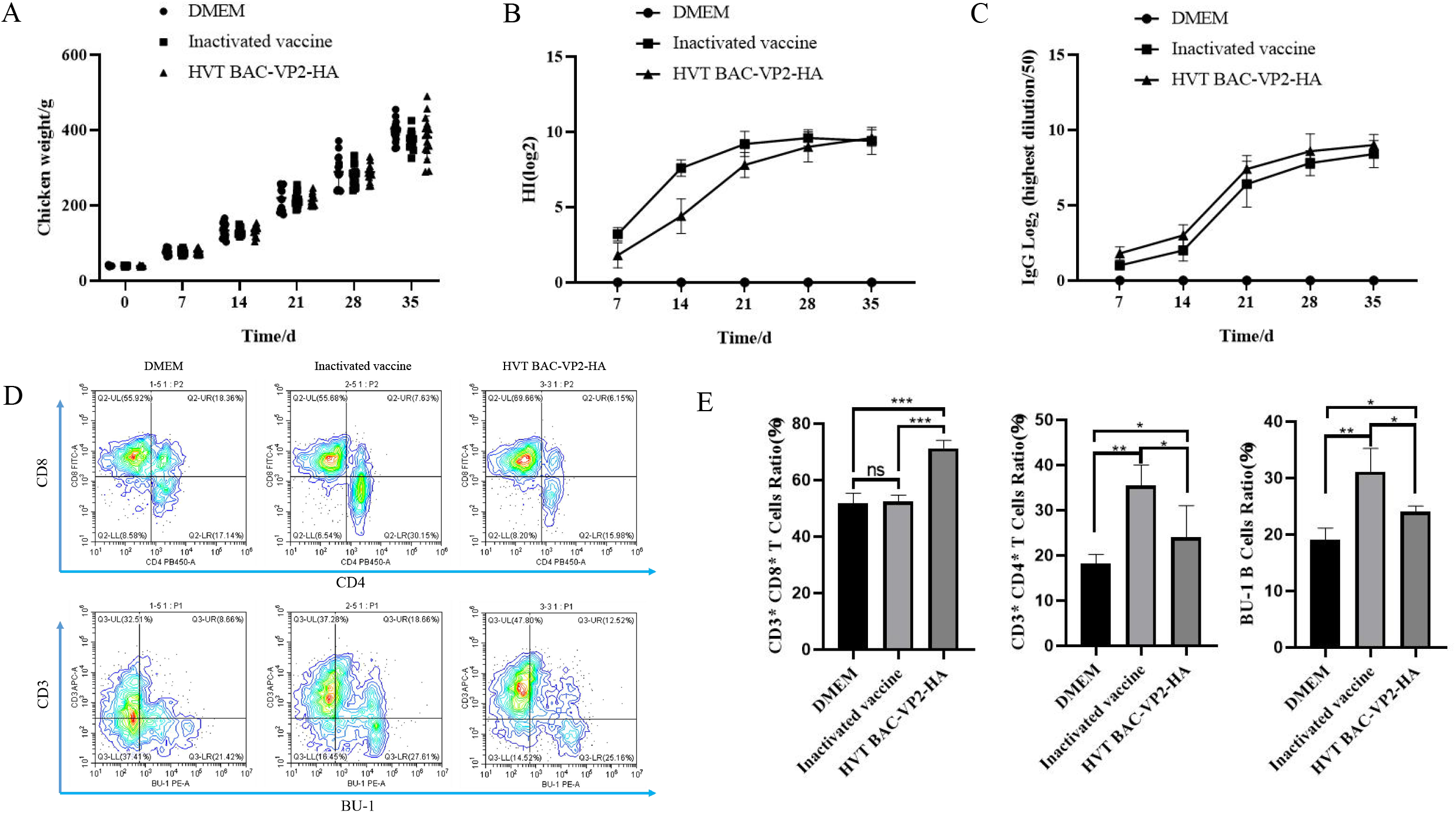
Evaluation of the safety and immunogenicity of the recombinant virus in SPF chickens. **(A)**. One-day-old SPF chickens were randomly divided into three groups and inoculated with DMEM (mock control), a commercial vaccine, or rHVT BAC-VP2-HA, respectively (n = 45). Body weights were measured at 7-day intervals for 35 days post-inoculation. **(B)**. Hemagglutination inhibition (HI) assay was used to determine HA-specific antibody titers in each group (n = 5). **(C)**. VP2-specific antibody levels were determined by enzyme-linked immunosorbent assay (ELISA) (n = 5). **(D)**. At 28 days post-vaccination, five chickens per group were randomly selected and euthanized. Splenic lymphocytes were isolated and analyzed by flow cytometry (n = 5). **(E)**. Frequencies of CD3^+^ T cells, CD3^+^CD4^+^ T cells, CD3^+^CD8^+^ T cells, and B cells in splenic lymphocytes were statistically analyzed (n = 5).

To evaluate the humoral immune responses induced by rHVT BAC-VP2-HA, blood samples were collected from five randomly selected birds per group at 7-day intervals post-immunization. HA-specific antibody titers were measured by HI assay, and VP2-specific antibody levels were determined by ELISA. Both rHVT BAC-VP2-HA and the inactivated vaccine induced robust HA-specific antibody responses, with titers progressively increasing, peaking at 28 days post-immunization, and subsequently reaching a stable plateau. No significant difference in HA-specific antibody levels was observed between the two vaccinated groups, whereas no detectable HI antibodies were found in the mock group (Fig. 3B). Similarly, VP2-specific ELISA revealed that both vaccines elicited strong antibody production, with comparable kinetics and peak levels at 28 days post-immunization. Again, no significant difference was detected between the two immunized groups, while VP2-specific antibodies were absent in the mock group (Fig. 3C).

To evaluate the cellular immune responses, five chickens per group were euthanized at 28 days post-vaccination, and splenic lymphocytes were isolated for flow cytometric analysis (Fig. 3D). The proportion of CD3^+^CD8^+^ T cells was significantly higher in the rHVT BAC-VP2-HA group than in both the mock and inactivated vaccine groups (*P* < 0.001), whereas no significant difference was observed between the mock and inactivated vaccine groups (Fig. 3E). In contrast, the proportions of CD3^+^CD4^+^ T cells and B cells were significantly higher in the inactivated vaccine group than in the rHVT BAC-VP2-HA group (*P* < 0.05), and both vaccinated groups showed higher levels than mock group (rHVT BAC-VP2-HA, *P* < 0.05; inactivated vaccine, *P* < 0.01; Fig. 3E). Collectively, these results indicate that rHVT BAC-VP2-HA elicits robust cellular immunity characterized by a strong CD8^+^ T-cell response.

### 3.4 rHVT BAC-VP2-HA confers effective protection against vIBDV challenge

To evaluate the protective efficacy of rHVT BAC-VP2-HA against vIBDV, 15 chickens from each of the rHVT BAC-VP2-HA group, inactivated vaccine, and the DMEM mock groups were challenged at 28 days post-vaccination, and additional five unchallenged chickens from the DMEM mock group serves as healthy controls.

Following challenge, the DMEM-challenge group exhibited typical clinical signs of IBDV infection, including depression, diarrhea, and anal pecking, accompanied by notable mortality. In contrast, no clinical symptoms or mortality were observed in either the rHVT BAC-VP2-HA or inactivated vaccine groups, indicating complete protection against vIBDV-induced morbidity and mortality.

To assess pathological damage, bursal tissues were collected at 7 dpc. Severe histopathological lesions, characterized by indistinct corticomedullary boundaries and extensive lymphocyte degeneration and necrosis, were observed in the DMEM-challenge group. By contrast, no obvious histopathological lesions were observed in either vaccinated group (Fig. 4A).

**Figure 4.**
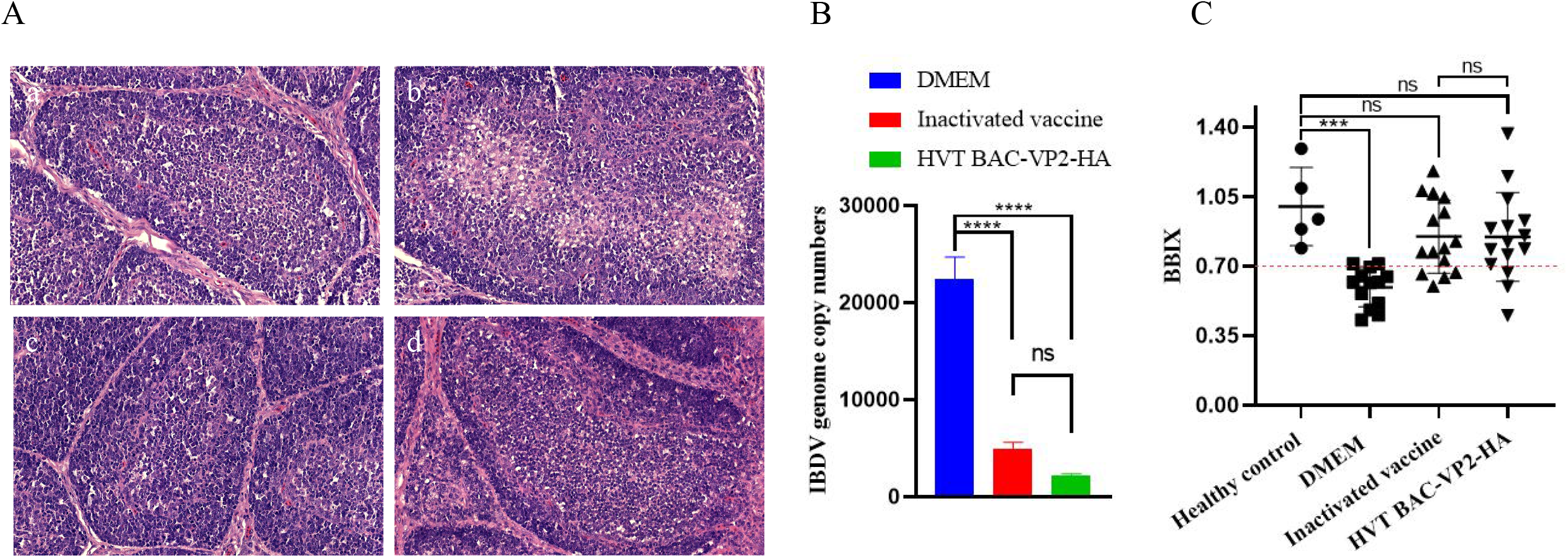
Histopathological lesions, viral loads, and bursal atrophy in SPF chickens following vIBDV challenge. **(A)**. Histopathological detection of the SPF chicken bursa after vIBDV challenge. (a) Healthy control group. (b) DMEM-challenge group. (c) Inactivated vaccine group. (d) rHVT BAC-VP2-HA group. **(B)**. Determination of viral loads in bursa of Fabricius following IBDV challenge. Total RNA was extracted from bursal tissues, reverse transcribed into cDNA, and viral genome copies were quantified by qPCR. **(C)**. Evaluation of bursal atrophy in SPF chickens following IBDV challenge. At 7 days post-challenge, all chickens were euthanized and necropsied. Bursa of Fabricius were collected and weighed, and the bursa-to-body weight ratio was calculated.

Viral loads in the bursa were determined to evaluate viral replication following challenge. Both vaccinated groups exhibited significantly lower viral loads than the DMEM-challenge group (*P* < 0.0001), with no significant difference between the rHVT BAC-VP2-HA and inactivated vaccine groups (*P* > 0.05; Fig. 4B).

Bursal atrophy was further assessed using the BBIX. No atrophy was observed in the healthy control group, whereas all chickens in the DMEM-challenged group exhibited marked atrophy (*P* < 0.001). Notably, both the rHVT BAC-VP2-HA and inactivated vaccine groups showed BBIX values comparable to those of the healthy control group, with no significant differences among them (*P* > 0.05; Fig. 4C).

Collectively, these results demonstrate that vaccination with rHVT BAC-VP2-HA confers robust protection against vIBDV challenge, as evidenced by the absence of clinical disease and mortality, a significant reduction in bursal viral replication, and effective prevention of bursal damage. The protective efficacy of rHVT BAC-VP2-HA is comparable to that of the commercial inactivated vaccine.

### 3.5 rHVT BAC-VP2-HA confers complete protection against H9N2 AIV challenge

To evaluate the protective efficacy of rHVT BAC-VP2-HA against H9N2 subtype AIV, chickens were challenged as described above and monitored for 7 days. No mortality was observed in any group throughout the observation period. However, chickens in the DMEM-challenged group began exhibiting clinical signs, including depression and diarrhea, at 3 dpc. In contrast, no clinical signs were observed in either the rHVT BAC-VP2-HA or inactivated vaccine groups during the entire observation period.

To assess viral shedding, oropharyngeal and cloacal swabs were collected at 3, 5, and 7 days post-challenge. All chickens in the DMEM-challenged group tested positive for viral shedding. In the inactivated vaccine group, only 1 out of 15 chickens was positive, yielding a protection rate of 93.3% (14/15). Remarkably, no viral shedding was detected in any chicken from the rHVT BAC-VP2-HA group, indicating complete protection (100%) (Fig. 5A). At 7 dpc, tracheal samples were collected for histopathological examination and viral load analysis. The DMEM-challenged group exhibited severe pathological lesions, including tracheitis, inflammatory cell infiltration, and epithelial necrosis with desquamation (Fig. 5B). In contrast, no AIV-associated pathological lesions were observed in either the rHVT BAC-VP2-HA or inactivated vaccine groups, resembling the healthy control group (Fig. 5B). Consistently, viral loads in the trachea were significantly reduced in both vaccinated groups compared with the DMEM-challenged group (*P* < 0.01; Fig. 5C).

**Figure 5.**
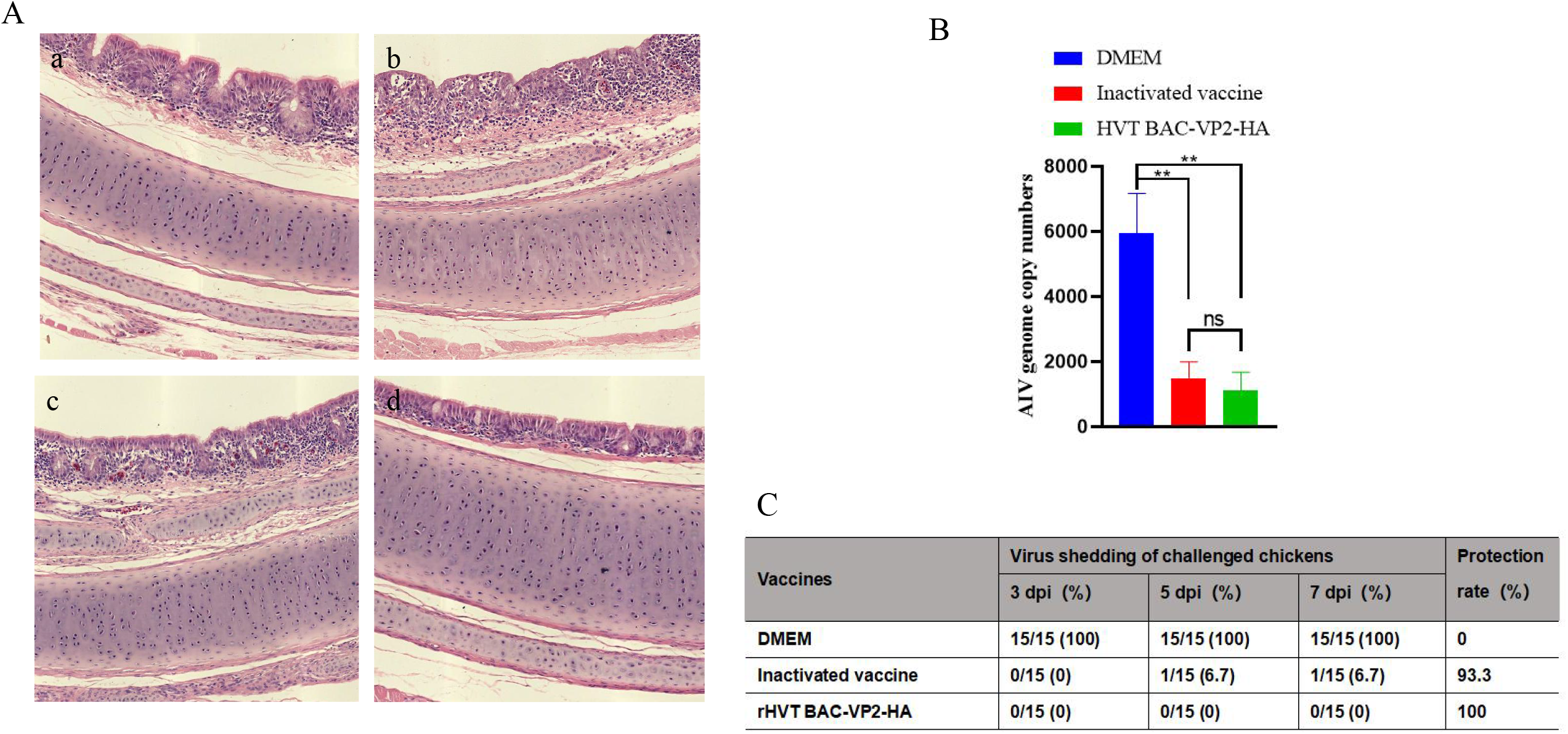
Histopathological lesions, viral loads, and virus shedding in SPF chickens following H9N2 AIV challenge. **(A)**. Histopathological detection of the SPF chicken trachea after H9N2 AIV challenge. (a) Healthy control group. (b) DMEM-challenge group. (c) Inactivated vaccine group. (d) rHVT BAC-VP2-HA group. **(B)**. Quantification of viral loads in tracheal tissues following H9N2 AIV challenge. Total RNA was extracted from tracheal tissues, reverse transcribed into cDNA, and viral genome copies were quantified by qPCR. **(C)**. Virus shedding and protection rates in chickens following H9N2 AIV challenge. Virus shedding was evaluated in oropharyngeal and cloacal swabs collected at 3, 5, and 7 days post-challenge and expressed as the number of positive chickens/total chickens (percentage). Protection rates were calculated based on the absence of detectable virus shedding.

Overall, rHVT BAC-VP2-HA vaccination not only prevented clinical disease but also completely blocked viral shedding following H9N2 AIV challenge, demonstrating superior protective efficacy compared with the inactivated vaccine under the conditions of this study.

## 4. Discussion

IBDV and H9N2 subtype AIV remain two major pathogens threatening the global poultry industry (Fan et al., 2019; Yu et al., 2026; Zhang et al., 2026a). IBDV infection causes severe immunosuppression, compromising the efficacy of subsequent vaccinations, whereas H9N2 AIV is widely prevalent and associated with persistent viral shedding and production losses (Dong et al., 2022; Fan et al., 2025; Kim et al., 2026). Although inactivated vaccines are commonly used to control these pathogens, they typically require multiple administrations and predominantly induce humoral immune responses, with limited ability to stimulate cellular immunity (Dong et al., 2022; Sagong et al., 2023). In contrast, recombinant HVT-vectored vaccines have emerged as a promising platform due to their ability to establish persistent infection, exhibit cell-associated viral spread, and continuously express foreign antigens (Li et al., 2016; van Hulten et al., 2021; Yang et al., 2025). This cell-associated infection pattern facilitates intracellular antigen processing and presentation via major histocompatibility complex class I (MHC-I) pathways, thereby promoting robust CD8^+^ T cell responses in addition to humoral immunity (Blander, 2023; Yamamoto et al., 2020). In the present study, we developed a recombinant HVT expressing both IBDV VP2 and H9N2 AIV HA proteins and systematically evaluated its biological characteristics, immunogenicity, and protective efficacy.

The successful construction and rescue of rHVT BAC-VP2-HA were confirmed by PCR, sequencing, and IFA, demonstrating stable integration and correct expression of both VP2 and HA (Fig. 1A-F). The *VP2* and *HA* genes were inserted into the *UL55* (HVT065/HVT066) and *RR* (HVT046/HVT047) loci, respectively. The recombinant virus exhibited plaque morphology and multistep growth kinetics comparable to those of the parental virus, indicating that insertion of foreign genes at these loci did not affect viral replication (Fig. 2A-B). Moreover, stable expression of *VP2* and *HA* was maintained over 20 serial passages, confirming the high genetic stability of the recombinant virus (Fig. 2C-D).

Immunization with rHVT BAC-VP2-HA did not adversely affect the growth performance of SPF chickens, confirming its safety *in vivo* (Fig. 3A). The recombinant vaccine induced robust VP2- and HA-specific antibody responses, with kinetics and magnitudes comparable to those induced by the commercial inactivated vaccine (Fig. 3B-C). Notably, however, rHVT BAC-VP2-HA elicited a significantly higher proportion of CD3^+^CD8^+^ T cells than the inactivated vaccine, whereas the latter induced relatively stronger CD4^+^ T cell and B cell responses (Fig. 3D-E). This distinct immune profiles likely reflect differences in antigen presentation pathways triggered by two vaccine types. As a live viral vector, HVT enables intracellular expression of foreign antigens, facilitating MHC-I-restricted antigen presentation and subsequent activation of cytotoxic T lymphocytes. In contrast, inactivated vaccines primarily induce humoral responses through exogenous antigen presentation. The enhanced CD8^+^ T cell response observed in the rHVT BAC-VP2-HA group may contribute to more efficient viral clearance following challenge.

Following challenge with vIBDV, rHVT BAC-VP2-HA provided effective protection, as evidenced by the absence of clinical signs and mortality, preservation of bursal structure, and a significant reduction in bursal viral load (Fig. 4A-C). The protective efficacy of rHVT BAC-VP2-HA was comparable to that of the commercial inactivated vaccine. Although no statistically significant difference was observed between the two vaccinated groups, the recombinant HVT vaccine offers additional advantages as a multivalent platform capable of simultaneously targeting multiple pathogens. Notably, complete immunity against IBDV is difficult to achieve, likely due to the virus’s tropism for the bursa of Fabricius and its immunosuppressive effects on the host, which may limit the development of full protective immunity even in vaccinated chickens. Consistent with this, mild bursal atrophy was observed in a small number of chickens in both vaccinated groups following challenge (Fig. 4C). Although the difference was not statistically significant, the lower incidence of bursal atrophy in the rHVT BAC-VP2-HA group compared with the inactivated vaccine group may suggest a trend toward improved preservation of bursal integrity.

In contrast, rHVT BAC-VP2-HA conferred complete protection against H9N2 AIV challenge, as indicated by the absence of clinical signs, a lack of viral shedding, and a significant reduction in viral loads in the respiratory tract (Fig. 5A-C). Importantly, no viral shedding was detected in the rHVT BAC-VP2-HA group, whereas a proportion of chickens in the inactivated vaccine group remained positive (Fig. 5C). This superior protective efficacy may be attributed to sustained antigen expression mediated by the HVT vector, which promotes long-term immune stimulation and enhances cellular immune responses, particularly CD8^+^ T cell-mediated viral clearance. CD8^+^ T cells play a critical role in antiviral immunity by eliminating infected cells, limiting viral replication, and reducing viral shedding. Furthermore, the use of viral endogenous promoters to drive foreign gene expression may contribute to more stable and sustained antigen expression, particularly *in vivo*, although their transcriptional strength is generally lower than that of strong heterologous promoters such as CMV.

## 5. Conclusion

Taken together, the results of this study demonstrate that rHVT BAC-VP2-HA is a safe and effective bivalent vaccine candidate that provides robust protection against both vIBDV and H9N2 AIV challenge. Notably, its ability to induce strong cellular immune responses and completely prevent viral shedding highlights its potential advantages over conventional inactivated vaccines. These findings support the further development and application of HVT-vectored multivalent vaccines for the control of major poultry diseases.

## Acknowledgments

This work was supported by the National Natural Science Foundation of China (grant number U21A20260), Natural Science Foundation of Henan Province (grant number 262300421479), the Key Scientific Research Program of 2025 for Colleges and Universities in Henan Province (grant number 25A230004).

## Author contributions

Yuanyuan Zhang: Data curation, Formal analysis, Investigation, Methodology, Writing - original draft

Xiuwen Yang: Data curation, Formal analysis, Investigation, Methodology, Writing - original draft

Yunzhe Kang: Data curation, Formal analysis, Investigation, Methodology, Writing - original draft

Wenhui Zhu: Formal analysis, Methodology

Yuanyuan Sun: Data curation, Methodology

Shaoyan Qi: Investigation, Methodology

Yuxin Chen: Investigation, Methodology

Guoqing Zhuang: Funding acquisition, Methodology, Project administration, Supervision, Writing - review and editing

Aijun Sun: Funding acquisition, Methodology, Project administration, Supervision, Writing - review and editing

## Data availability

All data are fully available without restriction.

## References

Alqazlan, N., Astill, J., Raj, S., Sharif, S., 2022. Strategies for enhancing immunity against avian influenza virus in chickens: a review. Avian Pathol 51, 211–235.

Blander, J.M., 2023. Different routes of MHC-I delivery to phagosomes and their consequences to CD8 T cell immunity. Semin Immunol 66, 101713.

Bruno, L., Nappo, M.A., Frontoso, R., Montinaro, S., Di Lecce, R., Guarnieri, C., Ferrari, L., Corradi, A., 2026. Avian Influenza Viruses: Global Panzootic, Host Range Expansion and Emerging One-Health Threats. Vet Sci 13.

Dong, J., Zhou, Y., Pu, J., Liu, L., 2022. Status and Challenges for Vaccination against Avian H9N2 Influenza Virus in China. Life (Basel) 12.

Fan, L., Wu, T., Hussain, A., Gao, Y., Zeng, X., Wang, Y., Gao, L., Li, K., Wang, Y., Liu, C., Cui, H., Pan, Q., Zhang, Y., Liu, Y., He, H., Wang, X., Qi, X., 2019. Novel variant strains of infectious bursal disease virus isolated in China. Veterinary Microbiology 230, 212–220.

Fan, W., Zeng, X., Chen, Y., Yu, Q., Zhang, Z., Tian, G., Liu, C., Bao, H., Qi, X., Wu, L., Zhang, Y., Liu, Y., Wang, S., Cui, H., Duan, Y., Chen, H., Gao, Y., 2025. A recombinant Marek’s disease vaccine candidate provides complete protection against infectious bursal disease virus and H9 subtype avian influenza virus in chickens. J Virol 99, e0114925.

Guo, X., Sun, W., Wei, L., Wang, X., Zou, Y., Zhang, Y., Li, S., Wang, N., Jiang, M., Zhao, H., Qu, E., Pang, Y., Yin, J., Ren, G., 2021. Development and evaluation of a recombinant VP2 neutralizing epitope antigen vaccine candidate for infectious bursal disease virus. Transbound Emerg Dis 68, 3658–3675.

Hein, R., Koopman, R., García, M., Armour, N., Dunn, J.R., Barbosa, T., Martinez, A., 2021. Review of Poultry Recombinant Vector Vaccines. Avian Dis 65, 438–452.

Ingrao, F., Rauw, F., van den Berg, T., Lambrecht, B., 2017. Characterization of two recombinant HVT-IBD vaccines by VP2 insert detection and cell-mediated immunity after vaccination of specific pathogen-free chickens. Avian Pathol 46, 289–299.

Jing, X., Tong, Q., Chen, W., Li, C., Jiang, Z., Sun, H., Sun, Y., Pu, J., Liu, J., Liu, L., 2026. Phylogenetic and pathogenic analyses of circulating infectious bursal disease virus strains in China. Poult Sci 105, 106108.

Kim, S.W., Park, J.Y., Son, J.E., Zheng, K.Q., Yu, C.D., Kim, K.W., Jeon, W.B., Choi, Y.R., Jang, H.K., Wei, B., Kang, M., 2026. Long-Term Immunogenicity and Protection of a rHVT-H9/Y280 Vaccine Against H9N2 Avian Influenza Virus in Commercial Layers with High Maternal Antibodies. Animals (Basel) 16.

Li, K., Liu, Y., Liu, C., Gao, L., Zhang, Y., Cui, H., Gao, Y., Qi, X., Zhong, L., Wang, X., 2016. Recombinant Marek’s disease virus type 1 provides full protection against very virulent Marek’s and infectious bursal disease viruses in chickens. Sci Rep 6, 39263.

Li, L., Shang, Y., Zhao, Q., Feng, H., Zeng, Z., Xiao, Q., Jiang, L., Yao, L., Wang, Z., Wang, H., Cheng, G., Luo, Q., Wen, G., 2026. Co-immunization with two recombinant Newcastle disease viruses expressing ILTV gB and H9N2 AIV HA confers protective efficacy against three avian pathogens. Poult Sci 105, 106661.

Liang, B., Fan, M., Meng, Q., Zhang, Y., Jin, J., Chen, N., Lu, Y., Jiang, C., Zhang, X., Zou, Z., Ping, J., Su, J., 2024. Effects of the Glycosylation of the HA Protein of H9N2 Subtype Avian Influenza Virus on the Pathogenicity in Mice and Antigenicity. Transbound Emerg Dis 2024, 6641285.

Liu, L., Wang, T., Wang, M., Tong, Q., Sun, Y., Pu, J., Sun, H., Liu, J., 2019. Recombinant turkey herpesvirus expressing H9 hemagglutinin providing protection against H9N2 avian influenza. Virology 529, 7–15.

Luczo, J.M., Spackman, E., 2024. Epitopes in the HA and NA of H5 and H7 avian influenza viruses that are important for antigenic drift. FEMS Microbiol Rev 48.

Sagong, M., Lee, K.N., Lee, E.K., Kang, H., Choi, Y.K., Lee, Y.J., 2023. Current situation and control strategies of H9N2 avian influenza in South Korea. J Vet Sci 24, e5.

Shah, A.U., Wang, Z., Zheng, Y., Guo, R., Chen, S., Xu, M., Zhang, C., Liu, Y., Wang, J., 2022. Construction of a Novel Infectious Clone of Recombinant Herpesvirus of Turkey Fc-126 Expressing VP2 of IBDV. Vaccines (Basel) 10.

Spackman, E., Stephens, C.B., Pantin-Jackwood, M.J., 2018. The Effect of Infectious Bursal Disease Virus-Induced Immunosuppression on Vaccination Against Highly Pathogenic Avian Influenza Virus. Avian Dis 62, 36–44.

Sun, H., Li, H., Tong, Q., Han, Q., Liu, J., Yu, H., Song, H., Qi, J., Li, J., Yang, J., Lan, R., Deng, G., Chang, H., Qu, Y., Pu, J., Sun, Y., Lan, Y., Wang, D., Shi, Y., Liu, W.J., Chang, K.-C., Gao, G.F., Liu, J., 2023. Airborne transmission of human-isolated avian H3N8 influenza virus between ferrets. Cell 186, 4074-4084.e4011.

Tischer, B.K., Kaufer, B.B., 2012. Viral bacterial artificial chromosomes: generation, mutagenesis, and removal of mini-F sequences. J Biomed Biotechnol 2012, 472537.

Tischer, B.K., von Einem, J., Kaufer, B., Osterrieder, N., 2006. Two-step red-mediated recombination for versatile high-efficiency markerless DNA manipulation in Escherichia coli. Biotechniques 40, 191–197.

van Hulten, M.C.W., Cruz-Coy, J., Gergen, L., Pouwels, H., Ten Dam, G.B., Verstegen, I., de Groof, A., Morsey, M., Tarpey, I., 2021. Efficacy of a turkey herpesvirus double construct vaccine (HVT-ND-IBD) against challenge with different strains of Newcastle disease, infectious bursal disease and Marek’s disease viruses. Avian Pathol 50, 18–30.

Wang, W., Wu, J., Jiang, N., Liang, Q., Liu, R., Fu, Q., Fu, G., Wei, T., Wan, C., Cheng, L., Huang, Y., He, X., Wei, P., Chen, H., 2025. Advances in Infectious Bursal Disease Virus Vaccines-A Review. Microorganisms 13.

Wang, Y., Jiang, N., Fan, L., Niu, X., Zhang, W., Huang, M., Gao, L., Li, K., Gao, Y., Liu, C., Cui, H., Liu, A., Pan, Q., Zhang, Y., Wang, X., Qi, X., 2021. Identification and Pathogenicity Evaluation of a Novel Reassortant Infectious Bursal Disease Virus (Genotype A2dB3). Viruses 13.

Wu, Z., Lin, X., Song, C., Feng, K., Ke, H., Yin, L., Fu, J., Yan, Z., Lin, W., Zhang, X., Chen, W., Xie, Q., 2026. Safety and immunogenicity of a broad-spectrum HVT vector vaccine for avian infectious bursal disease. Poult Sci 105, 105926.

Xie, Z., Chen, Y., Xie, J., Du, S., Chen, R., Zheng, Y., You, B., Feng, M., Liao, M., Dai, M., 2025. Construction with recombinant epitope-expressing baculovirus enhances protective effects of inactivated H9N2 vaccine against heterologous virus. Vet Microbiol 300, 110337.

Xiong, H., Wu, J., Xie, Q., Li, T., Wan, Z., Qin, A., Ye, J., Shao, H., 2025. Q221K mutation in VP2 drives antigenic shift of infectious bursal disease virus. Front Immunol 16, 1600371.

Yamamoto, K., Venida, A., Yano, J., Biancur, D.E., Kakiuchi, M., Gupta, S., Sohn, A.S.W., Mukhopadhyay, S., Lin, E.Y., Parker, S.J., Banh, R.S., Paulo, J.A., Wen, K.W., Debnath, J., Kim, G.E., Mancias, J.D., Fearon, D.T., Perera, R.M., Kimmelman, A.C., 2020. Autophagy promotes immune evasion of pancreatic cancer by degrading MHC-I. Nature 581, 100–105.

Yang, W., Liu, X., Wang, X., 2023. The immune system of chicken and its response to H9N2 avian influenza virus. Vet Q 43, 1–14.

Yang, W., Zhang, J., Dai, J., Guo, M., Lu, X., Gao, R., Liu, K., Gu, M., Hu, S., Liu, X., Wang, X., Liu, X., 2025. Multiple pathways to evaluate the immunoprotective effect of Turkeys Herpesvirus recombinant vaccine expressing HA of H9N2. Poult Sci 104, 104335.

Yu, H., Wang, G., Zhang, W., Wu, Z., Niu, X., Huang, M., Zhang, Y., Liu, R., Han, J., Xu, M., Han, J., Ling, D., Ke, E., Wang, S., Cui, H., Zhang, Y., Chen, Y., Liu, Y., Duan, Y., Gao, Y., Qi, X., 2025. Epidemiological characteristics of infectious bursal disease virus (IBDV) in China from 2023 to 2024: Mutated very virulent IBDV (mvvIBDV) is associated with atypical IBD. Poult Sci 104, 105195.

Yu, H., Wu, Z., Ling, D., Wang, G., Zhang, Y., Liu, R., Ke, E., Liu, X., Xu, T., Wang, S., Cui, H., Zhang, Y., Chen, Y., Liu, Y., Duan, Y., Gao, Y., Qi, X., 2026. First detection of natural co-infection with mutated very virulent IBDV (mvvIBDV) and novel variant IBDV (nVarIBDV). Poultry Science 105, 106365.

Zai, X., Shi, B., Shao, H., Qian, K., Ye, J., Yao, Y., Nair, V., Qin, A., 2022. Recombinant Turkey Herpesvirus Expressing H9N2 HA Gene at the HVT005/006 Site Induces Better Protection Than That at the HVT029/031 Site. Viruses 14.

Zhang, W., Sun, L., Wang, Q., Xia, C., Zhao, Y., 2026a. Molecular Characterization and Pathogenicity of H9N2 Avian Influenza Viruses in Poultry in Shandong Province, China, 2021–2023. Poultry Science, 106936.

Zhang, W., Wang, X., Gao, Y., Qi, X., 2022. The Over-40-Years-Epidemic of Infectious Bursal Disease Virus in China. Viruses 14.

Zhang, W., Wang, Y., Wang, G., Yu, H., Huang, M., Zhang, Y., Liu, R., Wang, S., Cui, H., Zhang, Y., Chen, Y., Gao, Y., Qi, X., 2025. Development and Application of Indirect ELISA for IBDV VP2 Antibodies Detection in Poultry. Viruses 17.

Zhang, Y., Yu, H., Wu, Z., Wang, G., Liu, R., Ling, D., Ke, E., Wang, S., Chen, Y., Liu, Y., Cui, H., Zhang, Y., Gao, Y., Qi, X., 2026b. Immune protection in chickens via lipid nanoparticle-encapsulated VP2 DNA vaccine against very virulent infectious bursal disease virus. Poult Sci 105, 106830.

